## Supplementary figures and images for "Biosynthetic gene cluster profiling predicts the positive association between antagonism and phylogeny in *Bacillus*"

### Supplementary Figure S1

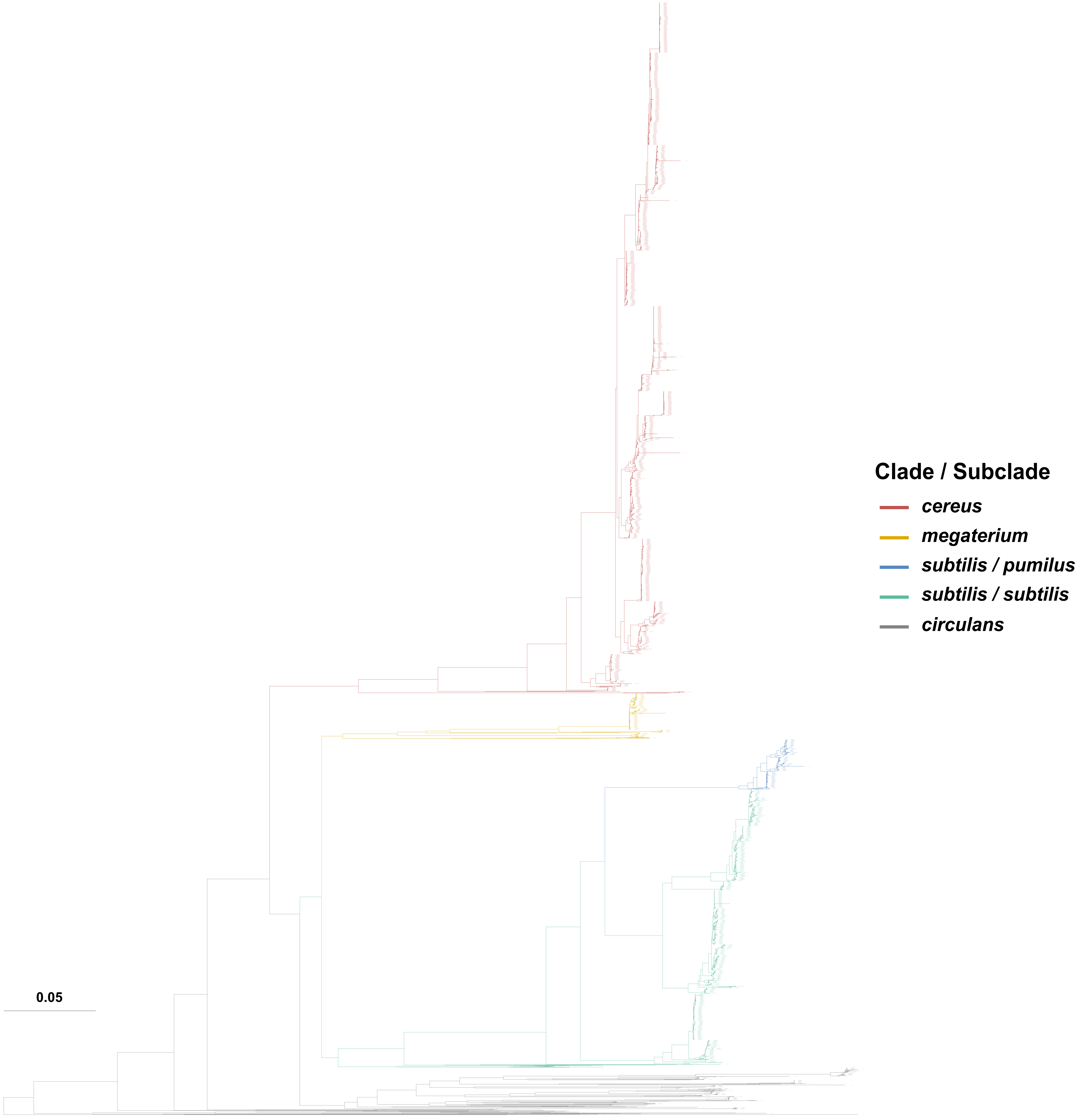

### Supplementary Figure S2

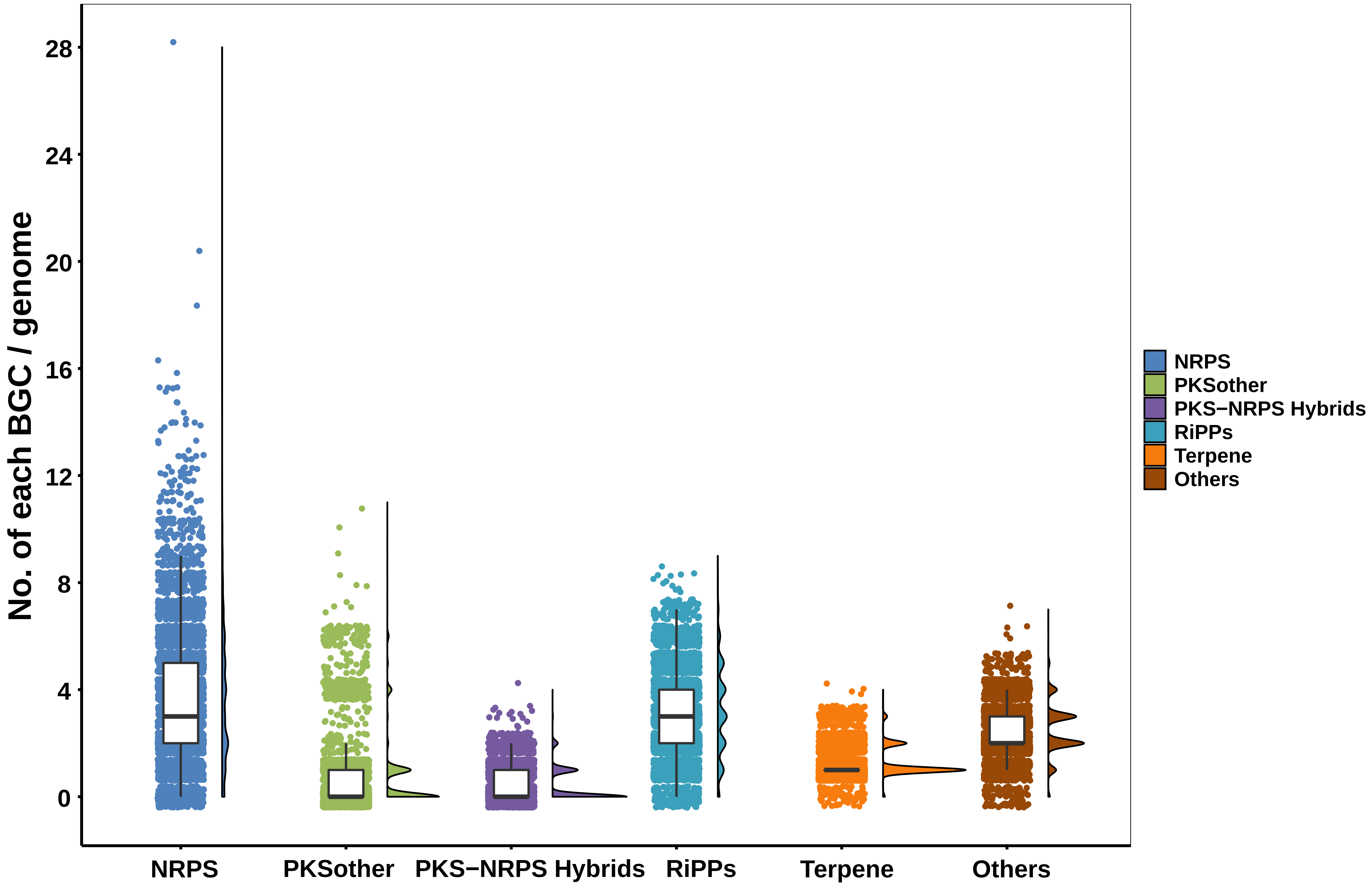

### Supplementary Figure S3

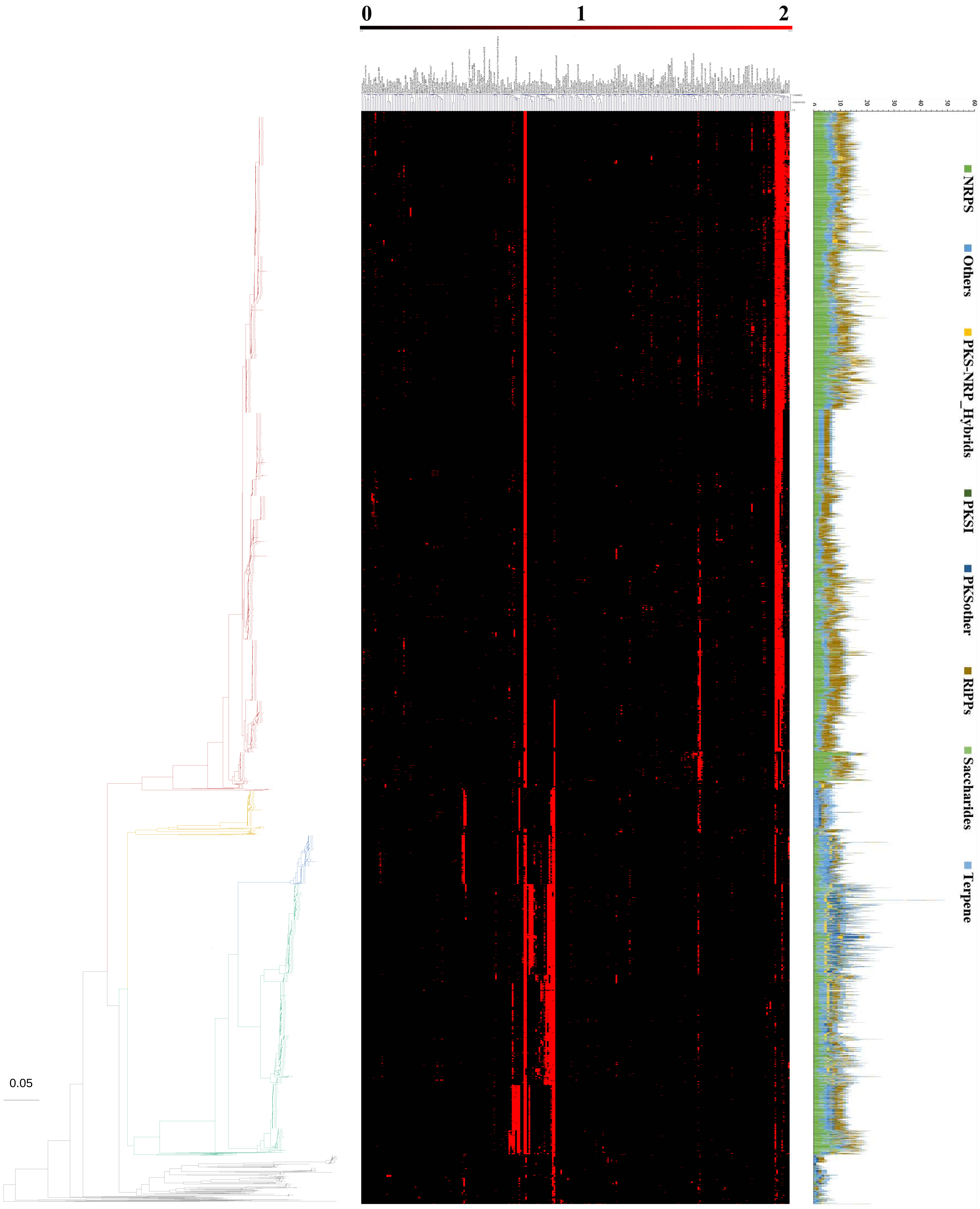

### Supplementary Figure S4

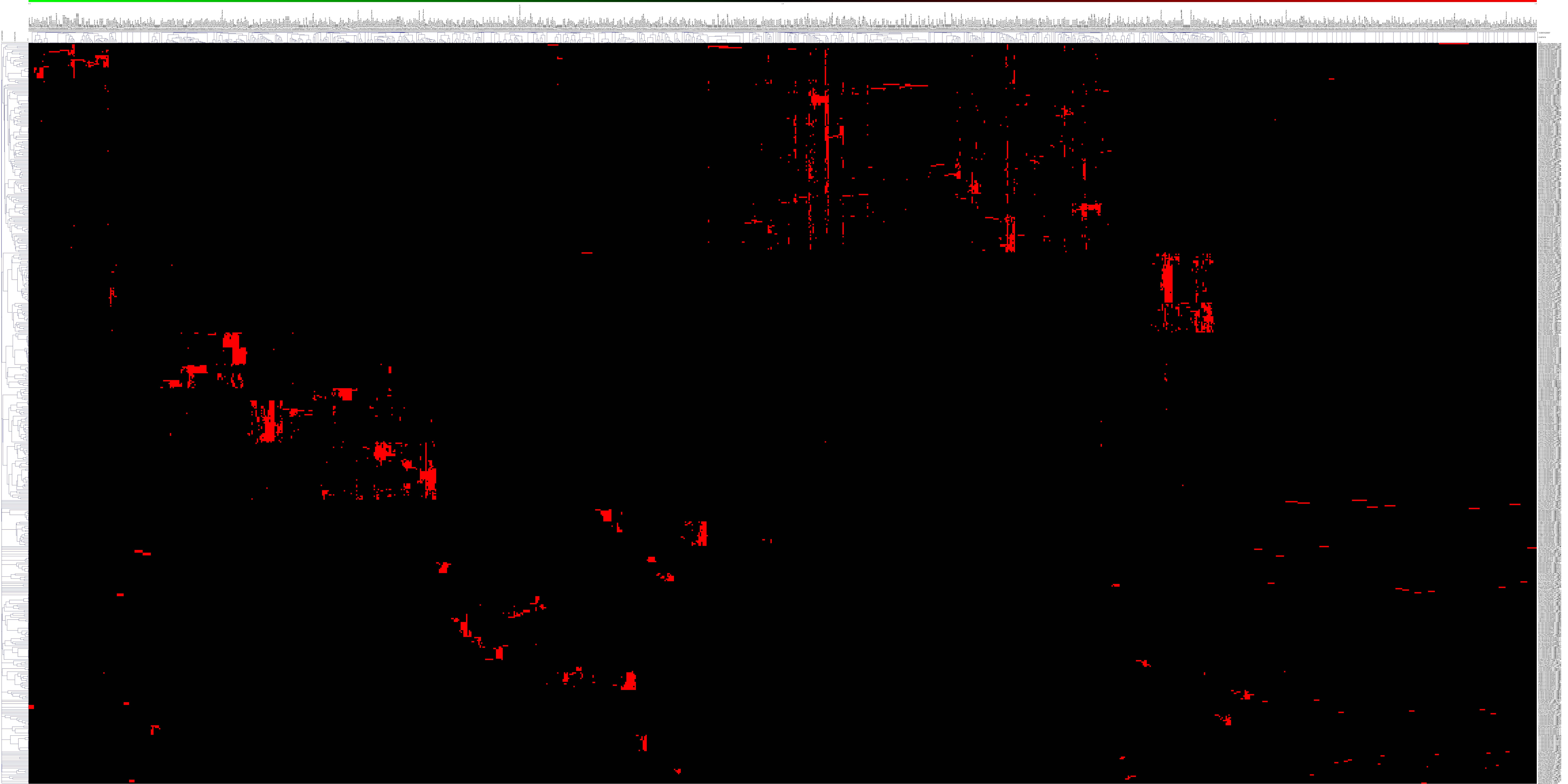

### Supplementary Figure S5

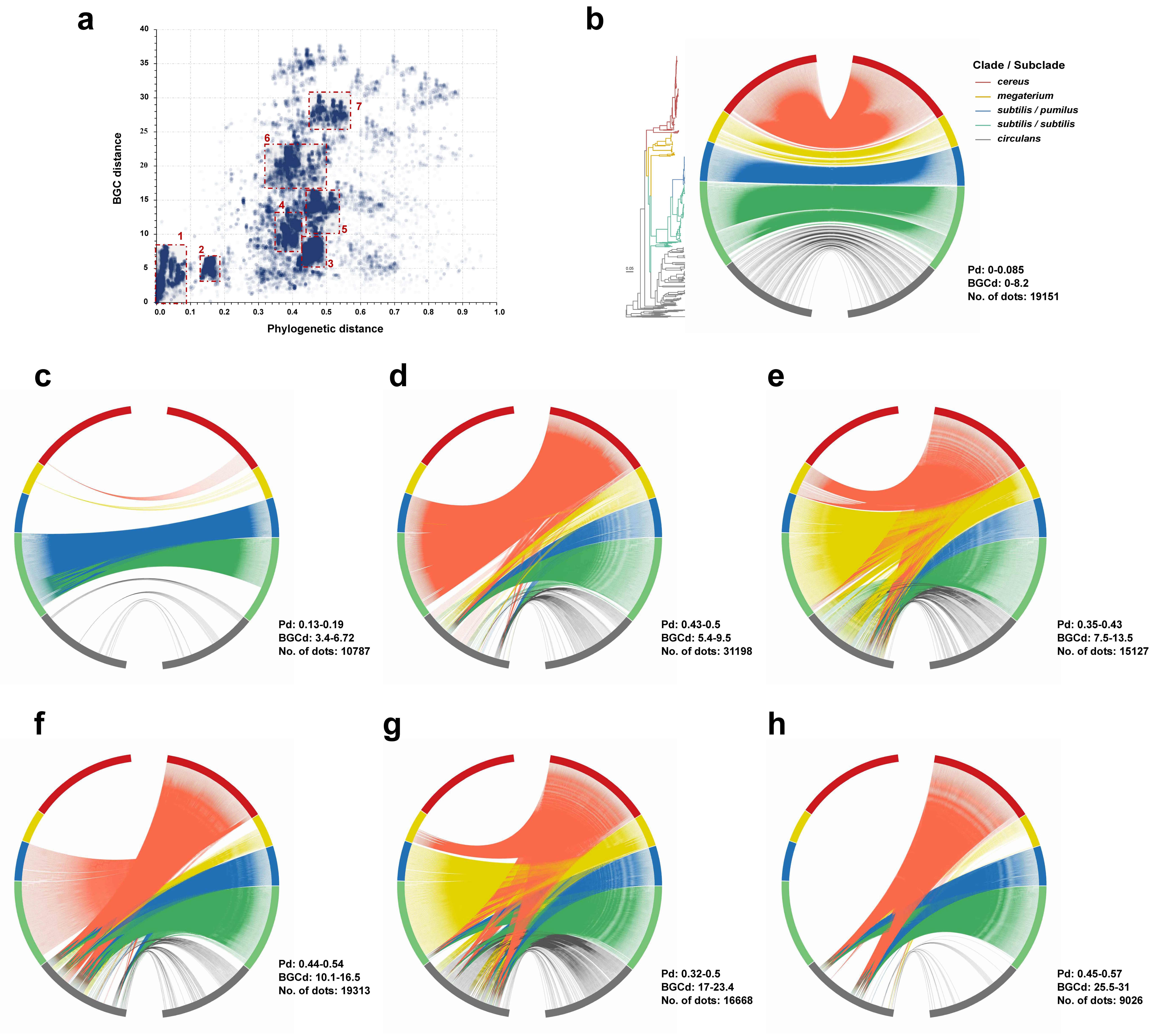

### Supplementary Figure S7

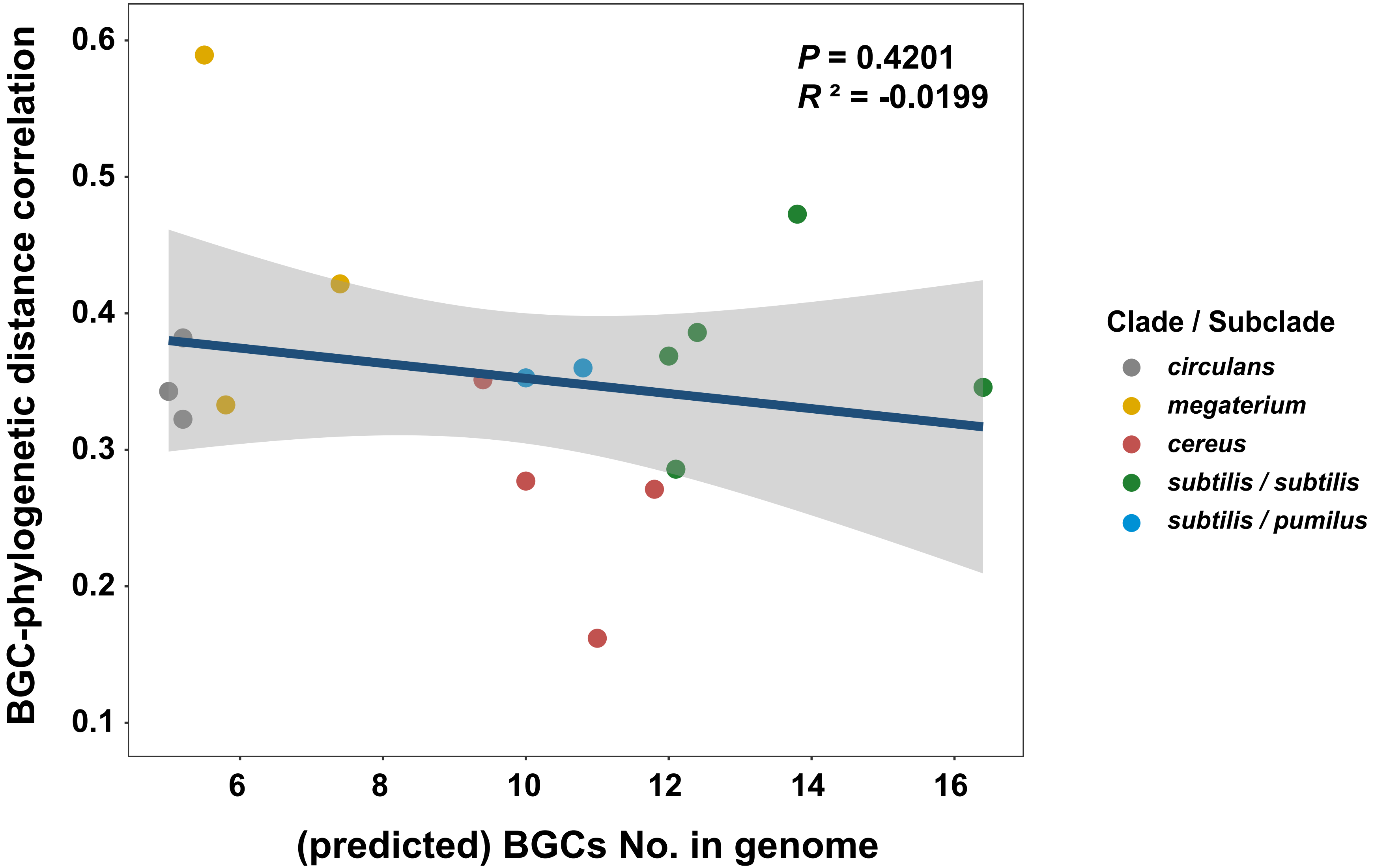
