## Supplementary Figure S6 for "Biosynthetic gene cluster profiling predicts the positive association between antagonism and phylogeny in *Bacillus*"

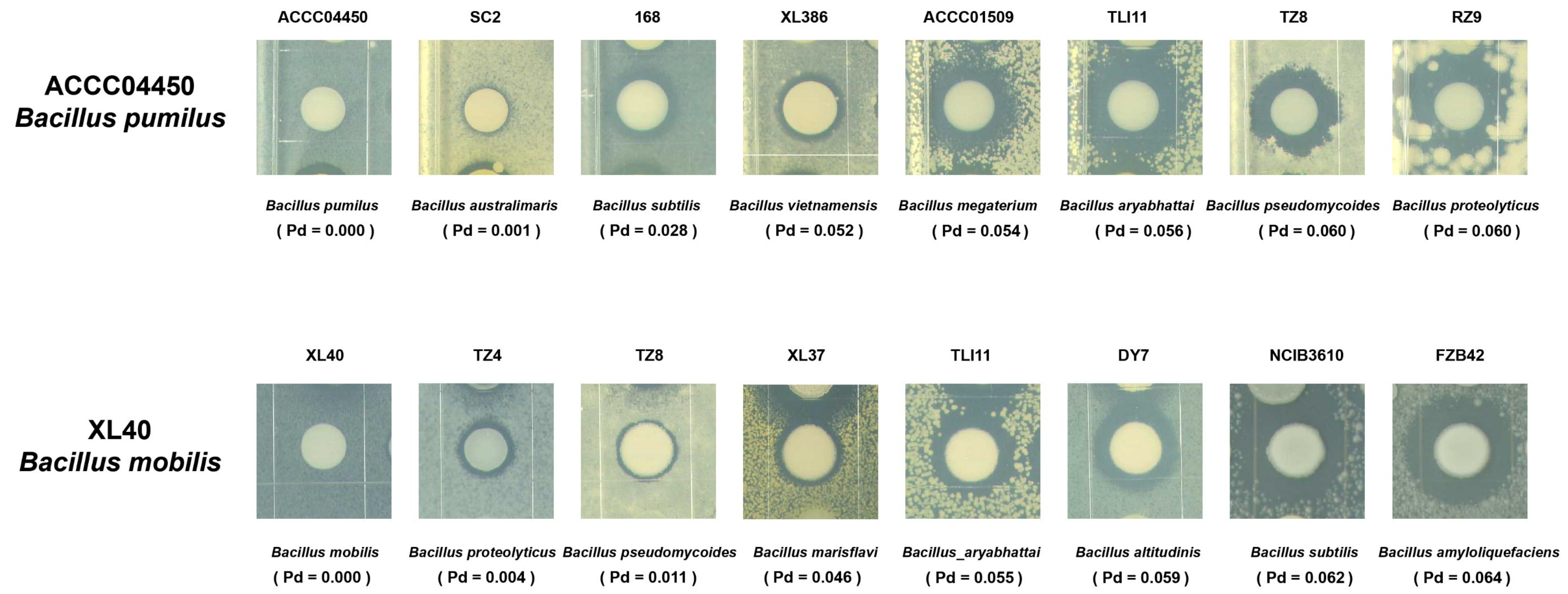


**Colony confrontation assay**

Pd: Phylogenetic distance

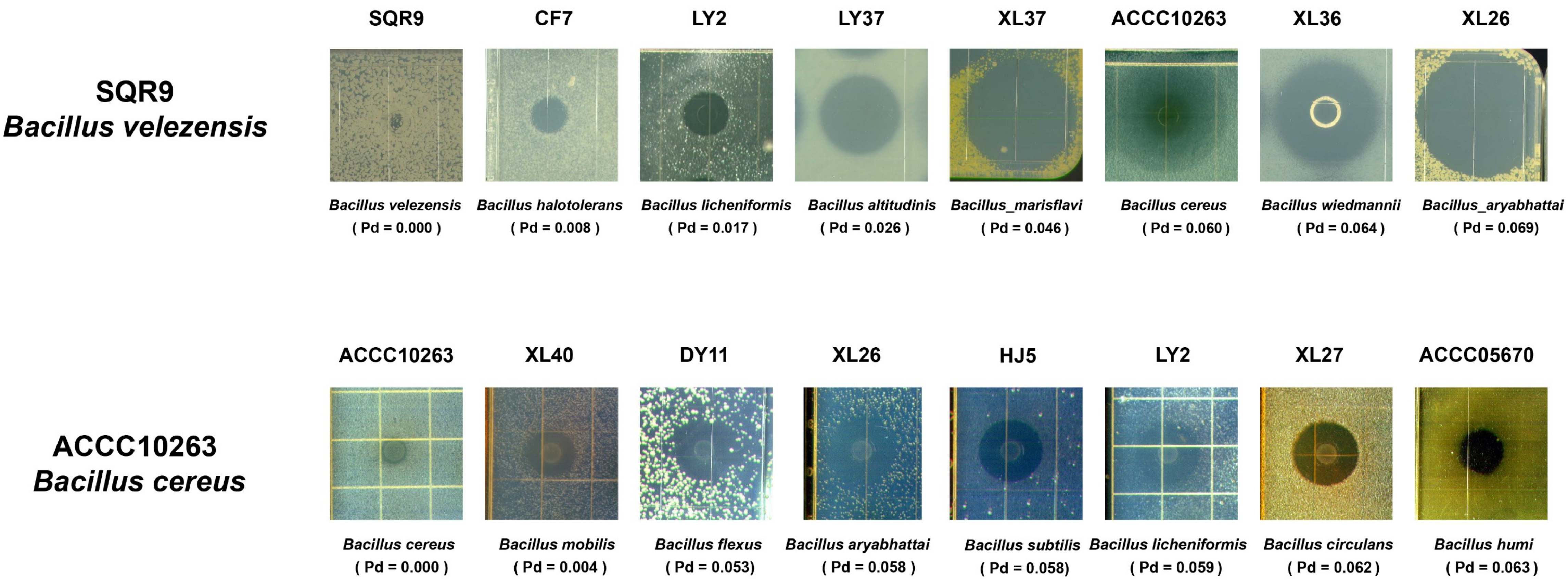


**Fermentation supernatant inhibition assay**

Pd: Phylogenetic distance
